# Rate variation and recurrent sequence errors in pandemic-scale phylogenetics

**DOI:** 10.1101/2024.07.12.603240

**Authors:** Nicola De Maio, Myrthe Willemsen, Zihao Guo, Abhratanu Saha, Martin Hunt, Nhan Ly-Trong, Bui Quang Minh, Zamin Iqbal, Nick Goldman

## Abstract

Phylogenetic analyses of genome sequences from infectious pathogens reveal essential information regarding their evolution and transmission, as seen during the COVID-19 pandemic.

Recently developed pandemic-scale phylogenetic inference methods reduce the computational demand of phylogenetic reconstruction from genomic epidemiological datasets, allowing the analysis of millions of closely related genomes. However, widespread homoplasies, due to recurrent mutations and sequence errors, cause phylogenetic uncertainty and biases.

We present new algorithms and models to substantially improve the computational performance and accuracy of pandemic-scale phylogenetics. In particular, we account for, and identify, mutation rate variation and recurrent sequence errors. We reconstruct reliable and public sequence alignment and phylogenetic tree of *>* 2 million SARS-CoV-2 genomes encapsulating the evolutionary history and global spread of the virus up to February 2023.

## 1 Introduction

Genomic epidemiology has become a vital tool in regional, national and global health, as exemplified during the COVID-19 pandemic[1–4]. It is a priority for national and international research, and its role in the interpretation and control of transmission of diverse human pathogens is expected to increase in future. Genomic epidemiology can reveal details about infectious pathogen biology[5], evolution[6], transmission[7], and effectiveness of containment measures[3]. These inform downstream research, but also have immediate effects for policy makers, in vaccine manufacturing, and in other areas.

Analysis of genomic epidemiological data relies heavily on phylogenetics. However, well-established phylogenetic methods have mostly been developed for inter-species evolutionary biology. Consequently, phylogenetic analysis of genomic epidemiological data with these methods is particularly challenging[8, 9]. Part of this challenge is due to the unprecedented size of such datasets, in particular those of SARS-CoV-2 genomes, with currently more than 20 million SARS-CoV-2 genomes shared globally. With ongoing improvement and widespread adoption of genome sequencing technologies, the sizes of genomic epidemiology datasets are expected to further increase in the future. Recently developed pandemic-scale phylogenetic methods, such as UShER[10, 11] and MAPLE[12] address the problem of computational demand in large genomic epidemiology datasets by employing algorithms and statistical tools specifically developed for this type of data. However, other difficulties affecting the analysis of such data remain. Of these, homoplasies (apparently recurring nucleotide substitutions along the phylogenetic tree) significantly affect the accuracy and reliability of phylogenetic inference[8]. We identified two predominant factors contributing to widespread homoplasies in SARS-CoV-2: (a) highly mutable nucleotides and genome positions[13], and (b) recurrent sequence errors [14, 15]. Addressing these two phenomena is expected to lead to more reliable phylogenetic estimation.

a. Accounting for heterogeneity in evolutionary rates is common and consequential in phylogenetics[16]. In popular phylogenetic models of rate variation, each genome position is allowed to evolve at any of a given number of rates[17], which usually leads to more accurate inference[18]. These approaches, however, substantially increase computational demand[19] and might not be able to account for the extreme level of mutation rate variation observed in genomic epidemiological data sets[13]. To overcome these problems, here we propose a computationally efficient model in which each genome position is assigned its own estimated mutation rate, similar to the SLR model[20] of heterogeneous selection pressure. This approach leverages the large size of available datasets to model in detail highly variable mutation rates.
b. Recurrent sequence errors are ubiquitous in SARS-CoV-2 genomic data[14, 15], and are highly detrimental to phylogenetic estimation due to the low levels of sequence divergence in such data[15]. Here, we extend previously proposed phylogenetic sequence error models (which typically assume homogeneity in errors along the genome[21, 22]) to explicitly model and infer position-specific sequence error probabilities. This allows us to not only effectively identify positions with recurrent errors, which can then be masked from the sequence alignment[14, 15, 23], but also to account for these errors during phylogenetic inference itself.

We implemented these models within MAPLE, together with algorithmic improvements, to reduce the computational demand of the software. We use these tools, millions of publicly shared SARS-CoV-2 genomes, and recent advancements in consensus calling methods to reconstruct a publicly available, reliable and global SARS-COV-2 phylogenetic tree.

## 2 Results

### 2.1 New models and algorithms

Models and algorithms presented here are discussed in detail in the Methods Section 5 and have been implemented within our approximate maximum likelihood phylogenetic software MAPLE v0.6.8 https://github.com/NicolaDM/MAPLE.

#### 2.1.1 Models of rate variation and recurrent sequence errors

In addition to the GTR[24] substitution model, we consider three extensions:

1. The more general non-stationary, non-reversible UNREST nucleotide model [25], which can account for the non-stationary genome evolution of SARS-CoV-2 in human hosts[13]. Since MAPLE avoids matrix exponentiation[12], our estimation is not affected by numerical instability. Given the low divergence in the considered datasets, We approximate the root nucleotide frequencies with the observed nucleotide frequencies in the alignment.
2. Secondly, we extend the UNREST model to account for rate variation. Differently from classical random variable models in phylogenetics[17], our approach allows one rate per genome position, assigning a free parameter to each site to model its position-specific substitution rate. This approach is highly parameterized, and therefore only suitable for large genomic datasets, and allows us to account for elevated variation in substitution rates, and in particular highly mutable sites, which are a cause of extensive homoplasies in SARS-CoV-2[13, 15]. This model is described in detail in Section 5.2.
3. Lastly, we extend the rate variation UNREST model above to account for recurrent sequence errors. In this model each position of the genome is assigned a site-specific error probability free parameter. Again, this site-specific sequence error model is highly parameterized and is only suitable for large datasets. More details about this model are given in Section 5.3.

We only consider the rate variation model as an extension of the UNREST substitution model. Our rationale is that using a more restricted substitution model such as GTR in conjunction with rate variation would mean that differences in substitution rates across nucleotides that are not accounted for by the GTR model (e.g. elevated C to T and G to T mutation rates increasing T genome content over time[13]) might lead to wrongly infer rate variation (e.g. higher rates at C and G genome positions). Similarly, we only consider the error model as an extension of the rate variation model, that is, we never estimate error probabilities without also estimating rate variation. The reason is that, without accounting for rate variation, the error model might wrongly interpret excess mutations at a site as the result of high error probability.

We efficiently infer the values of the thousands of free parameters in our more complex models using Expectation Maximization (see Section 5.4). We give a graphical example in Fig. 1 to explain how we can estimate mutation rate and error probability jointly for every position of the genome. Without sequence errors, we expect substitutions to happen along the phylogenetic tree at the same rate for every branch, and so hit terminal and internal branches indiscriminately (e.g. Fig. 1B) and proportionally to their length. A recurrent sequence error instead is expected to cause an enrichment in the number of substitutions observed at terminal branches (e.g. Fig. 1C, D). We estimate site-specific substitution rates and error probabilities by distinguishing substitutions at internal vs terminal branches, while at the same time accounting for different substitution rates for different nucleotides, for branch lengths, and for uncertainty in the nucleotide substitution history.

**Figure 1:**
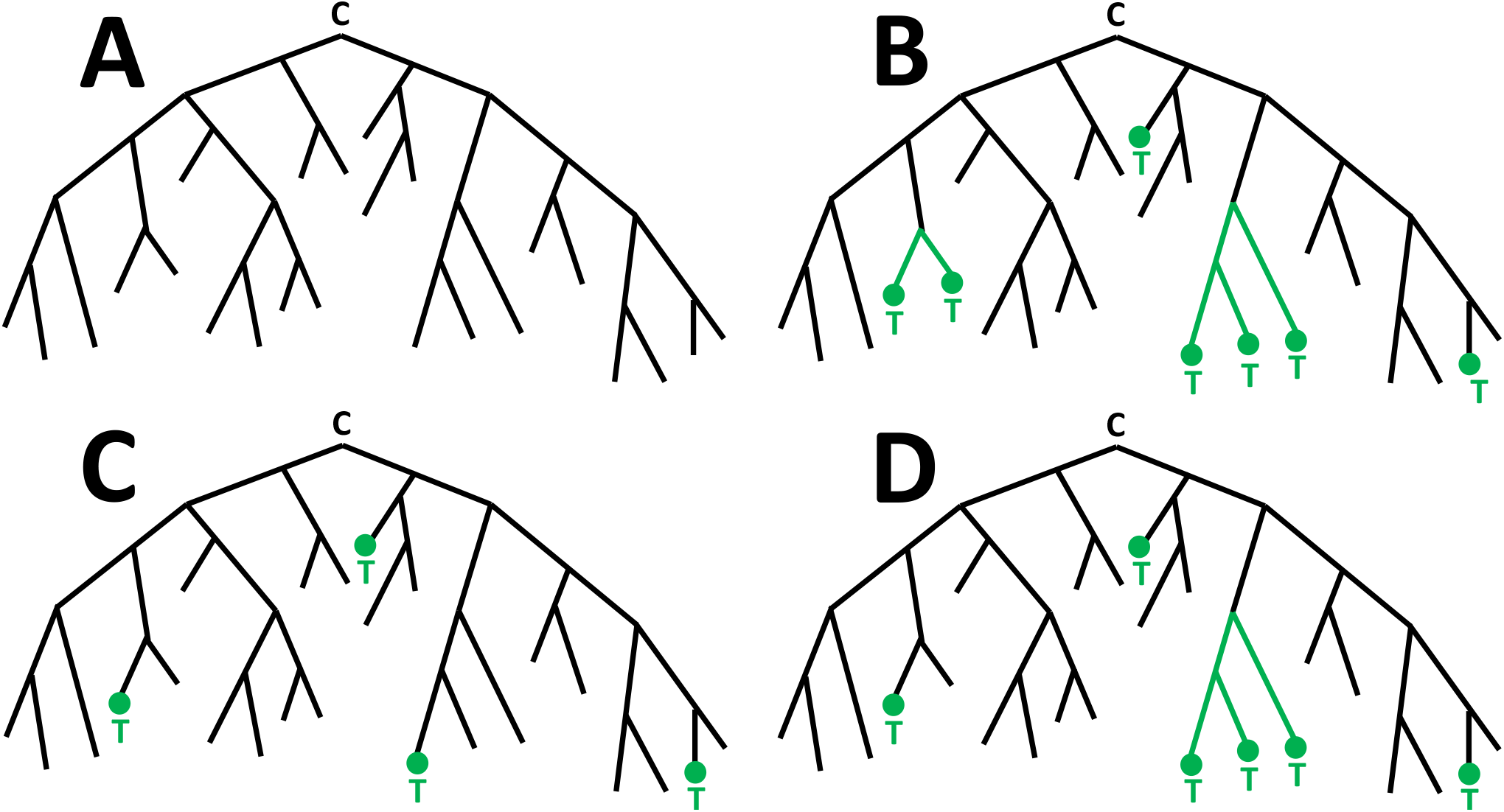
Graphical examples of the effects of substitution rate and error probability variation. In these examples nucleotide C is the root state, while nucleotide T (shown in green in the terminal nodes as well as on branches) is the derived nucleotide. For simplicity, we show a very small phylogeny. In practice, in SARS-COV-2 phylogenetic trees substitutions are more numerous and much sparser, and so there is less uncertainty in substitution history and parameter inference. **A** When mutation rate and error probability at a given genome position are both 0, the corresponding alignment column will have only one nucleotide and no substitutions. **B** The presence of mutations, but not sequence errors, is expected to result in a balanced proportion of observed substitutions in internal and terminal branches (here as an example we have 2 mutations on internal branches and 2 on terminal ones). **C** A site with no mutations (0 mutation rate) but positive error probability results in an enrichment of apparent substitutions at terminal branches (here we have 4 substitutions on terminal branches). **D** A site with positive mutation rate and error probability is expected to result in substitutions at internal branches, but also an enrichment of apparent substitutions at terminal branches (here one substitution at an internal branch and 3 at terminal ones).

#### 2.1.2 Local references

MAPLE performs phylogenetic analysis of vast genomic epidemiological data by representing closely related observed genome sequences, reconstructed ancestral genomes, and their uncertainty, in terms of differences with respect to a reference genome[12]. However, as an outbreak progresses, divergence with respect to the reference genome increases, and as such the computational demand of storing and processing genetic data under this paradigm also increases. In principle, information could be stored more efficiently: a mutation-annotated tree[26] can represent observed genomic sequences efficiently by recording estimated mutation events on the tree. This means that an observed genome can be reconstructed by tracing, along a phylogenetic tree, the inferred mutational history separating this genome from the root, ancestral genome. This way, each mutation event only needs to be represented once in the phylogenetic tree, and does not need to be explicitly represented in the descendants of the branch where the mutation occurred.

The problem with using a mutation-annotated tree in phylogenetics is, however, that information for two distant nodes of the tree cannot be directly compared. For any comparison, one needs to cross the branches in the tree separating the two considered nodes and update their information (‘translating’ it so that both are represented based on the same background reference) accordingly. We found that while this approach leads to a reduction in memory consumption, it did not substantially reduce runtime. Instead, our algorithm selects a sparse number of tree nodes and uses them as local references (Fig. 2). A useful reference is not too similar to another reference, and is used to represent sufficiently many genomes, so we select nodes based on thresholds on the number of descendant genomes and the number of mutations separating it from its most recent reference ancestor. Genetic sequence information and uncertainty at nodes in the tree is then represented in terms of their closest reference ancestor. The advantage of this approach is that, when comparing information across different nodes in the tree, only a small number of ‘translations’ needs to be carried out (one for each traversed local reference), while substantial computational demand is saved thanks to the more concise representation of genetic sequence information.

**Figure 2:**
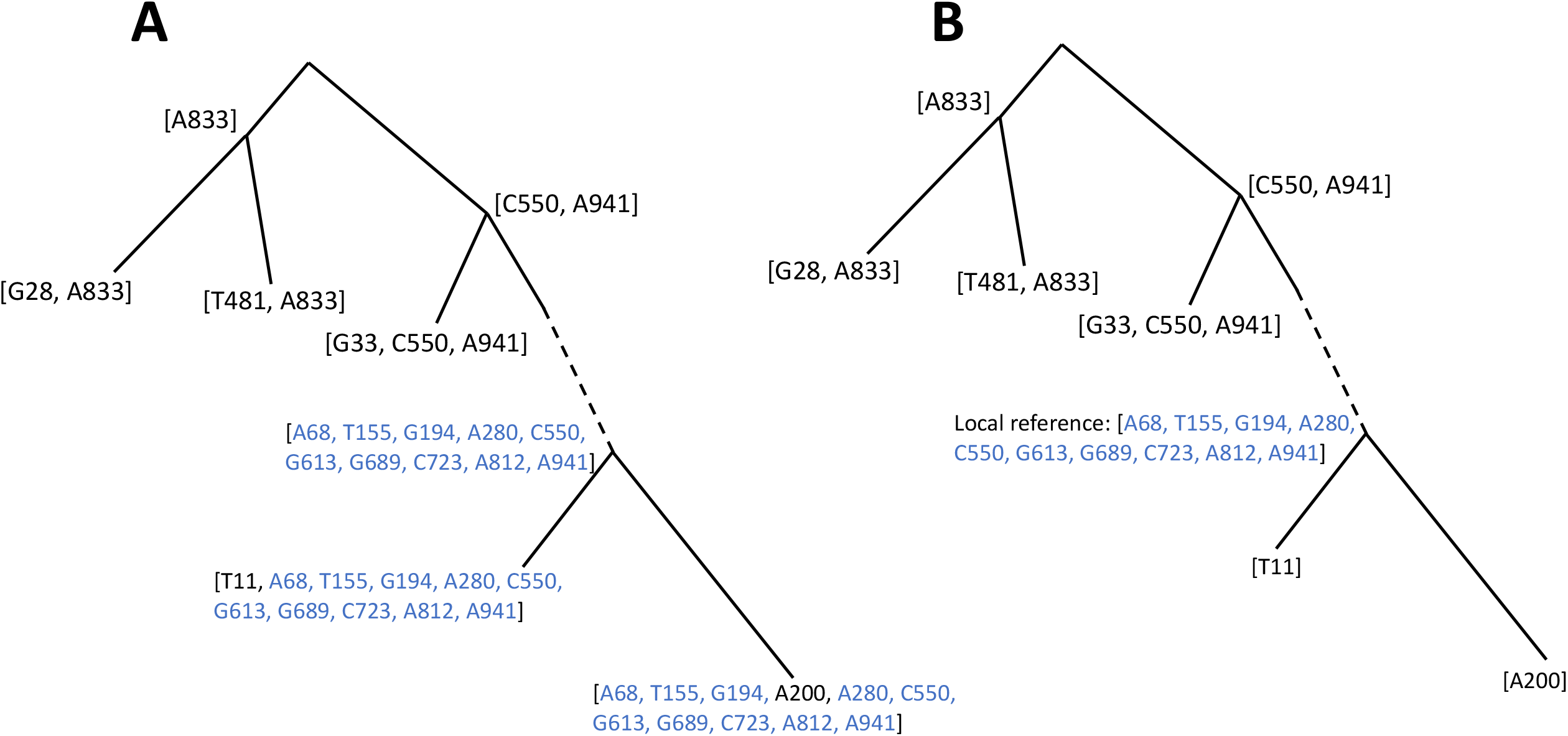
Graphical representation of local references. Previously, we represented observed and ancestral genomes in terms of differences with respect to a fixed reference genome[12], as exemplified in **A**. Here, for graphical simplicity, the ancestral genome of the root of the tree is assumed to be the reference genome. We highlight in blue the mutations accumulated along the dashed branch which in this case are observed and represented in all the descendants of this branch. Our new representation of the same tree and genome evolution history is showcased in **B**. Here a new local reference is defined at the bottom of the dashed branch; the genome of a local reference is represented in terms of its differences with respect to its parent reference (in this case the root genome). All genomes downstream of a local reference (except those that are eventually downstream of a further local reference) are represented in terms of their differences with respect to this local reference. This makes the representation of these genomes more concise, and their comparison faster.

#### 2.1.3 Parallelization

The most time- and memory-intensive part of the MAPLE algorithm is the topological improvement search through SPR moves[12]. We have now parallelized this step to speed up the analysis of large collections of genomes. MAPLE can now partition nodes/subtrees of an initial phylogenetic tree into non-overlapping sets, and assign each set to a different core. Each core then performs SPR searches for re-placement of its assigned subtrees in parallel. When all cores have finished their search, the possible SPR moves found for each node/subtree re-placement are ranked based on the likelihood improvement they are expected to bring to the tree. Starting from the most promising node/subtree to be re-placed, a new serial SPR search is then performed, with the best found SPR moves being this time actually implemented by modifying the tree before moving to the next initially most promising node/subtree to be re-placed. Because the number of advantageous SPR moves found is typically small compared to the total number of nodes in the tree, this final serial step is typically very fast compared to the initial parallelized SPR search.

#### 2.1.4 Other new features

In the Methods Section we present additional improvements we made to MAPLE, such as allowing fast and accurate estimation of branch lengths (Section 5.5.4), improved topological search (Section 5.5.2), efficient tree root search (Section 5.5.3), and allowing sample placement onto an initial tree for online phylogenetic inference (Section 5.5.1).

### 2.2 Computational demand

Our improvements lead to approximately halving of time and memory demand of MAPLE (Table 1). Runtime can further be reduced by parallelizing the topological search, but since this is not currently done in a shared memory framework, it also leads to an increase in memory demand (Table 1). Also, expectedly, attempting to parallelize with too many cores leads to a reduction in performance.

**Table 1:**
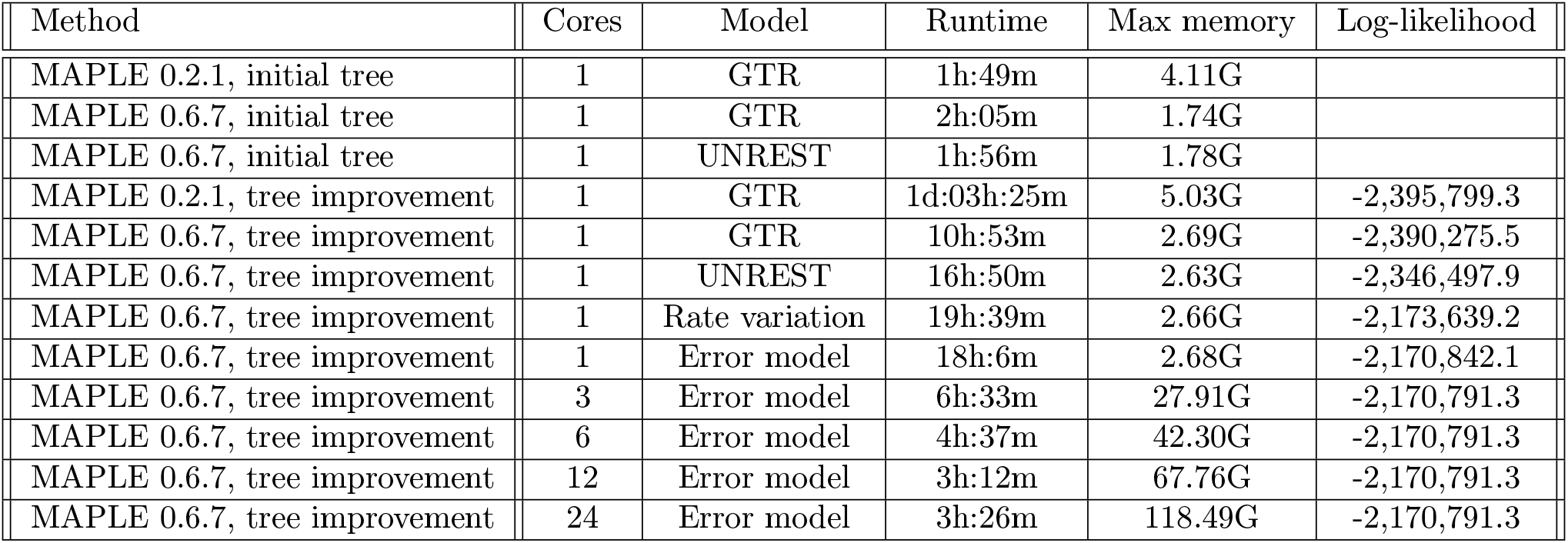
Performance of different algorithms and models analysing a 100,000 SARS-CoV-2 genomes subsample of the real Viridian dataset (see Section 5.6.1). We separate initial tree inference (first 3 rows) from tree improvement (rest of the table) since the latter can be parallelized in MAPLE 0.6.7, while the former cannot. Tree improvement in MAPLE 0.2.1 was started from the MAPLE 0.2.1 initial tree. Tree improvement in MAPLE 0.6.7 with GTR model was started from the initial tree from MAPLE 0.6.7 with GTR model. All other tree improvement runs were started from the initial tree from MAPLE 0.6.7 with UNREST model.

**Table 2:** Masked Viridian genome alignment positions. We also masked (not included in the table) position 25202 for computational convenience (it contains no alternative nucleotide calls, but 534,413 ambiguous calls), positions 22195, 22197, 22198, 22202, and 22204 due to recurrent alignment errors causing 5 linked artificial substitutions at these positions in about 50,000 genomes, and positions 28245, 28247, 28249, 28351, 28253 and 28254 for similar reasons. For the other positions we highlight in red what we consider red flags in the consensus alignment, read data, and the phylogenetic inference. ‘Minor variants’ means that *>* 5% frequency minor variants at these positions were systematically observed in the mapped reads of genomes inferred by MAPLE to contain a consensus sequence error. ‘Ambiguities’ means the number of ambiguous nucleotide characters counted in Viridian consensus sequences - higher numbers highlighted in red. ‘Consensus differences’ means differences in numbers of alternative nucleotides in Viridian vs GenBank consensus sequences; large differences are highlighted in red. ‘Phylogenetic clustering’ means phylogenetic clustering of errors and substitutions at the considered positions, assessed by visually inspecting the estimated phylogenetic tree annotated with inferred errors and substitutions. ‘Substitution rate’ refers to the substitution rate estimated by MAPLE; the genome-wide average rate is 1.0, high rates are highlighted in red. ‘Likely errors’ is our guess of the number of Viridian consensus sequence errors at the considered position, after taking all evidence into account. ‘Notes’ contains additional information, such as apparent or known causes of the errors at these positions. For these preliminary analyses, positions 1-72 and 29769-29903 were preventively masked due to being not called in a large fraction of the genomes.

| Position | Minor variants | Ambiguities | Consensus differences | Phylogenetic clustering | Substitution rate | Likely errors | Notes |
|---|---|---|---|---|---|---|---|
| 76 |  | 154 | 30 As vs 93 |  | 0.5 | < 100 | Low coverage |
| 78 |  | 18 | 10 As, 15 Cs, 187 Gs vs 80 As, 30 Cs, 51 Gs |  | 1.5 | < 200 | Low coverage |
| 274 | X | 391 | 355 As vs 9 | X | 1.4 | ≈ 300 | Low coverage CA mutations |
| 4321 | X | 628 | 707716 Ts vs 709324 | X | 4.6 | Hundreds | Heterozygous reversions in BA.2 |
| 8008 |  | 239 | 20135 As vs 2 | X | 26 | ≈ 20000 | Low coverage, correlated with 8012 |
| 8012 |  | 10 | 4678 Cs vs 1 | X | 13 | ≈ 4600 | Low coverage, correlated with 8008 |
| 8826 |  | 77 | 131 Cs, 237 Gs vs 31 Cs, 1 G | X | 7.2 | < 400 | Low coverage, correlated with 8829 |
| 8829 |  | 301 | 801 As vs 33 | X | 1.6 | Hundreds | Low coverage, correlated with 8826 |
| 8835 | X | 52213 | 357989 Cs vs 68802 | X | 100 | ≈ 360000 | ARTIC4 alternative primer binding[14].<br>Correlated with 15521. |
| 15510 |  | 713 | 1916 Cs vs 733 | X | 34 | ≈ 1900 | Assembly difficulties |
| 15521 | X | 31294 | 756265 As vs 188634 | X | 100 | ≈ 750000 | ARTIC4 alternative primer binding[14].<br>Correlated with 8835 |
| 15854 | X | 55 | 424 As vs 8 | X | 1.4 | ≈ 400 |  |
| 17259 | X | 141 | 7898 Gs vs 6677 | X | 15 | ≈ 7000 |  |
| 19413 | X | 1332 | 1532 Gs vs 74 | X | 32 | ≈ 1500 |  |
| 19672 | X | 199 | 365 As vs 182 | X | 0.09 | < 400 | Heterozygosity at 19668 |
| 21650 | X | 1110 | 95 Cs vs 4 | X | 0.7 | < 100 |  |
| 21987 | X | 34374 | 421580 Gs vs 451142 | X | 100 | Thousands | ARTIC3 amplicon dropout[29] |
| 22027 |  | 11 | 1548 Gs vs 11 | X | 13 | ≈ 1500 | Mapping issues near deletion 22029-34 |
| 22033 |  | 82 | 2364 Ts vs 712 | X | 1.3 | ≈ 1500 | Mapping issues near deletion 22029-34 |
| 22786 | X | 18127 | 643395 Cs vs 671746 | X | 100 | Thousands | Issues downstream AC mutation |
| 22882 | X | 1881 | 1069445 Gs vs 1109262 | X | 100 | Thousands | Issues downstream TG mutation |
| 23118 | X | 39690 | 528 Ts vs 68 | X | 1.6 | ≈ 500 |  |
| 23948 | X | 1257 | 1361584 Ts vs 1374120 | X | 25 | Thousands | Issues downstream GT mutation |
| 25296 | X | 888 | 177 Cs vs 9 | X | 2.2 | < 200 |  |
| 25324 | X | 22 | 418 As vs 0 | X | 0.7 | ≈ 400 | Correlated with 25336 |
| 25336 | X | 1212 | 413 As vs 572 | X | 1.9 | ≈ 400 | Correlated with 25324 |
| 26530 | X | 4098 | 653150 Gs vs 659866 | X | 22 | Thousands | Heterozygous reversions in BA.1 |
| 26766 | X | 4190 | 1688 Cs vs 2141 | X | 1.9 | ≈ 1700 |  |
| 27507 | X | 842 | 41636 Cs vs 44029 | X | 40 | Hundreds | Issues downstream AC mutation |
| 29687 |  | 10 | 251 Cs vs 69 | X | 3.6 | < 300 | Low coverage |

Since our rate variation model allows only one evolutionary rate per genome position, it leads to limited additional computational cost compared to the more basic models (Table 1 and Fig. 3A-B). Due to the low divergence of the considered datasets, our expectation maximization approach can estimate thousands of model parameters (such as site-specific rates and error probabilities) at limited additional time and memory demand. However, more complex models in MAPLE can still lead to some increase in runtime, for example due to the increased complexity of the tree search associated with the lowering of the likelihood cost of the most recurrent mutation events.

**Figure 3:**
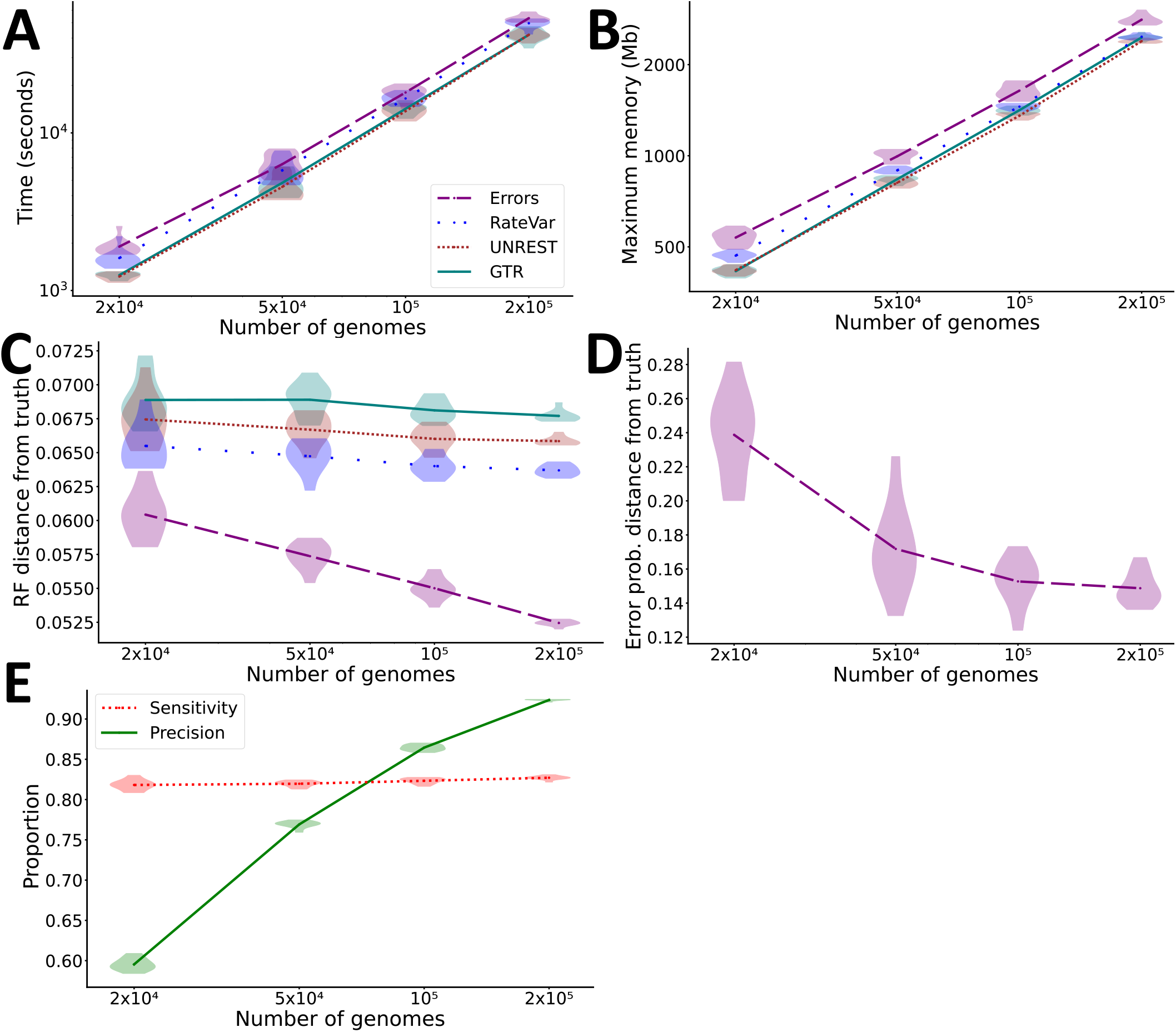
Simulation-based benchmark. Assessment of different models using SARS-CoV-2-like genome data simulated with rate variation and recurrent sequence errors (see Section 5.6.2). On the X axis are genome subset sizes. For each size and model considered we ran 10 replicates. Dots show the mean across replicates, while violin plots show variation between replicates. The four models considered are GTR, UNREST, UNREST with rate variation (‘RateVar’), and UNREST with rate variation and sequence errors (‘Errors’). **A** Time demand and **B** maximum memory demand for non-parallelized tree inference from scratch. **C** Normalized Robinson-Foulds distances between simulated (‘true’) phylogenetic trees and estimated ones; lower values represent more accurate estimations of tree topology. **D** Euclidean distance between the vectors of estimated and simulated position-specific sequence error rates; lower values represent more accurate estimates of position-specific error parameters. **E** Precision of inference of individual sequence errors (true positives divided by total inferred errors) and sensitivity (true positives divided by total simulated errors). We used a probability threshold of 50% to define inferred sequence errors within input genome sequences. The total number of simulated errors was ≈ 0.22 per genome. All analyses were run on one core of an Intel Xeon Gold 6252 Processor @ 2.10 GHz.

### 2.3 Simulation-based benchmark

To assess the accuracy of our methods, we aimed at simulating realistic SARS-CoV-2 genome data including both mutation rate variation and position-specific error probabilities (Section 5.6.2). Knowing the exact phylogenetic tree used in simulations and the position of sequence errors generated in the genome data, we can assess MAPLE’s accuracy in estimating both. We find that our more complex models, and in particular the error model, substantially improve phylogenetic tree inference accuracy (Figure 3C). MAPLE needs sufficiently large datasets to reliably identify position-specific error parameters and errors in the input genome sequences (Figure 3D-E), with accuracy increasing as more sequences are considered. With 200,000 genomes, we have approximately 92% precision and 83% sensitivity in estimating individual sequence errors.

### 2.4 Towards reliable, global, and public SARS-CoV-2 alignment and phylogeny

Currently, millions of SARS-CoV-2 genome sequencing read datasets are shared publicly. This represents a unique opportunity to reliably and consistently call millions of consensus genomes, and to corroborate putative recurrent consensus sequence errors.

To validate our methods, we collected millions of SARS-CoV-2 genome sequencing read datasets (see Section 5.6.1). First, we investigated highly recurrent sequence errors (Section 2.4.1). Then, we filtered out putative recurrent sequence errors and possibly contaminated samples to obtain a reliable, global, and public SARS-CoV-2 sequence alignment and phylogeny (Section 2.4.2).

#### 2.4.1 Investigation of recurrent sequence errors

We collected SARS-CoV-2 genomes with publicly available sequencing read data, a consensus sequence in GenBank, and a Viridian[27] consensus genome. We consider Viridian genomes here since they have been consistently assembled and called using an approach that addresses reference biases affecting consensus genomes from public databases[27]. After filtering out low-coverage genomes and samples that might have been affected by contamination or mixed infections, our collection contained 2,993,121 genomes (see Section 5.6.1). For these, we created two alignments, one containing the GenBank consensus genomes, and one containing the Viridian ones. Since we want to investigate the presence and abundance of recurrent sequence errors in these alignments, for now we do not perform any prior masking of alignment columns (e.g. [14, 23]). From each alignment we inferred a phylogeny and recurrent sequence errors with MAPLE. We then investigated read data and the phylogeny to corroborate candidate recurrent sequencing or bioinformatics artifacts in the consensus genomes. In particular, we focused on positions inferred to contain errors with a frequency around or above 0.01% of genomes. We considered the following red flags of possible recurrent artefacts:

1. The presence, at the considered sample and genome position, of multiple nucleotides supported at high frequency (*>* 5%) by reads. This pattern can be, for example, a result of primers binding at unintended sites of homology[14].
2. Elevated numbers of IUPAC ambiguity characters in the consensus sequences. In the presence of high frequency alternative nucleotides, consensus calling pipelines can include ambiguity characters instead of calling a nucleotide. Recurrent ambiguity characters are therefore a symptom of recurrent alternative nucleotides in the reads at the considered genome position.
3. Frequent different consensus sequence calls by Viridian and GenBank at the considered position. This suggests that different protocols for read mapping, filtering, and trimming, or consensus calling thresholds might systematically affect the consensus sequences at the considered position.
4. Phylogenetic clustering of apparent substitutions caused by putative errors. This can happen if errors are genotype-dependent (e.g. a real mutation might affect primer binding, or an indel might affect quality of read alignment to the reference genome), or if errors are caused by sequencing protocols whose adoption might vary by time and location. We visually investigated mutation-annotated phylogenies inferred by MAPLE with Taxonium [28].
5. High mutation rate estimated by MAPLE at the considered position. MAPLE’s error model assumptions can be substantially violated if the frequency of errors at a given position is too high (e.g. *>* 1%), if they are correlated with other errors, or if they are not uniformly distributed along the phylogenetic tree. In these cases, portions of the recurrent errors can be interpreted by MAPLE as genuine mutations, leading to high estimated mutation rates at positions affected by recurrent errors.

While analysing the GenBank consensus genome alignment, we noticed that many sites of high-frequency mutations (those with hundreds of thousands of descendants) had large numbers of artificial reversions to the reference genome, resulting from genomes with low coverage at these positions. Some of the sites with the most putative erroneous reversions in GenBank consensus genomes are positions 210, 7124, 9053, 21618, 22813, 23854, 26107, 26577, 27373, 28881, 28916, 29402, and 29742 of the reference genome MN908947.3. This appears to be the result of common reference biases in consensus calling pipelines, which are addressed by Viridian[27]. For this reason, we focused the rest of our analyses on the Viridian consensus sequences, and used these as the basis to create a reliable global SARS-CoV-2 alignment and phylogeny.

A few positions of the Viridian genomes were inferred to contain recurrent errors, particularly positions 8835 and 15521, which were previously also highlighted as problematic, since they are frequently affected by artifacts caused by ARTIC V4 primers binding to unintended genome locations[14, 23]. In Table 2.4.1 we list positions inferred to be error-prone in SARS-CoV-2 Viridian genomes and that we mask from the alignment.

Other positions inferred by MAPLE to contain moderate numbers of errors (in the order of hundreds) showed no red flags corroborating possible recurrent consensus sequence errors. We suspect that these might be caused by other phenomena resulting in an enrichment in the number of substitutions on terminal branches, for example deleterious recurrent mutations. The positions that seem most affected by this pattern are 4579, 8318, 11522, 13424, 21255, 21550, 22487, 23416, 25163, 25166, and 26029.

#### 2.4.2 Alignment masking and tree inference

With the aim to create a global, reliable SARS-CoV-2 alignment and phylogeny, we aligned all available Viridian SARS-C0V-2 genomes, masked genome ends and putative error-prone positions, and filtered out low-coverage and highly heterozygous samples which could be affected by contamination or mixed infection. See Section 5.6.1 for a detailed description of our data preparation. The final alignment contains 2,072,111 Viridian consensus genomes.

From this alignment, we inferred a phylogenetic tree using MAPLE under an UNREST substitution model with rate variation. Inference of the initial tree with one core took about 4 days 17 hours and 18.5G maximum memory; topological improvement search parallelized over 14 cores took about 5 days 13 hours and 440G maximum memory. The alignment, metadata, inferred tree, and inferred substitution rates have been uploaded on Zenodo [30].

Consistently with previous analyses[13], genome positions 10323, 11083, 21137, and 27384 are inferred to be those with the highest substitution rate (Figure 4). In particular, we infer *>* 11, 000 separate mutation events at position 11083 in our phylogenetic tree.

**Figure 4:**
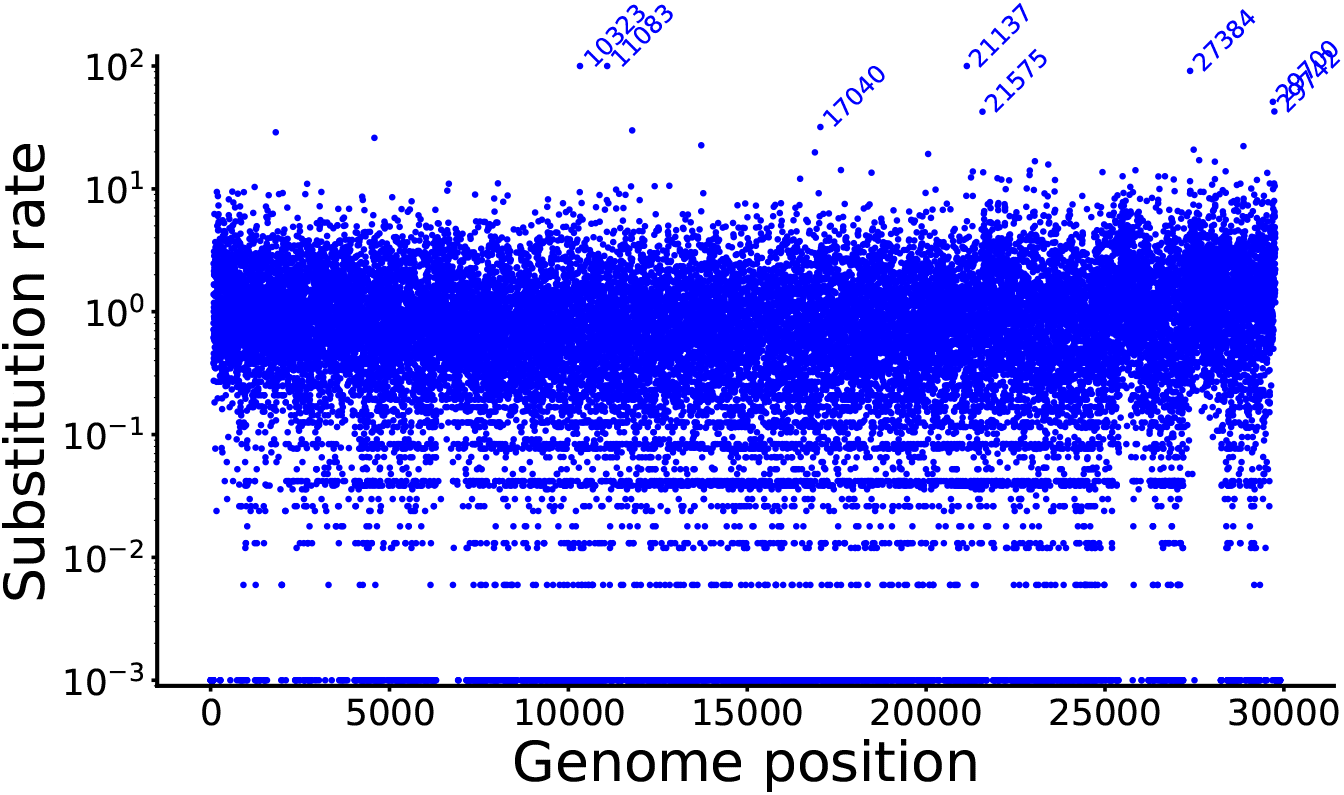
Estimated position-specific SARS-CoV-2 substitution rates. Site-specific values of the relative substitution rates (maximum 100 and minimum 0.001 allowed) estimated under the rate variation model in MAPLE from the masked and filtered real *>* 2*M* Viridian consensus real SARS-CoV-2 genome alignment. High values of substitution rate (*>* 30) are annotated with the corresponding genome position.

It is not simple to systematically compare our phylogenetic tree with other similar ones, such as the global, public, SARS-CoV-2 tree cov2tree[28, 31], due to differences in genomes included, consensus calling method, and alignment masking approach. While we see strong overlap in the two trees regarding inferred evolutionary history of major SARS-CoV-2 variants, we also notice some differences. For example, we infer one of the major Delta branches to be composed of two sister clades, the first one containing lineages AY.13, AY.14, and AY.54 among others, and the second, larger one, containing lineages AY.9 and AY.47 among many others (Figure 5). cov2tree instead infers the first clade to be a sub-clade of the second one. Investigating the cov2tree tree and mutational history, it appears that the latter represents a local tree optimum, and that removing and re-attaching the first clade in a similar way as in the MAPLE tree leads to a more parsimonious evolutionary history, with 2 fewer substitution events (Figure 5). This is exactly the kind of tree improvements that we have designed MAPLE to perform with its subtree prune and regraft (SPR) search, and this example shows the positive impact it has for inferring phylogenetic trees.

**Figure 5:**
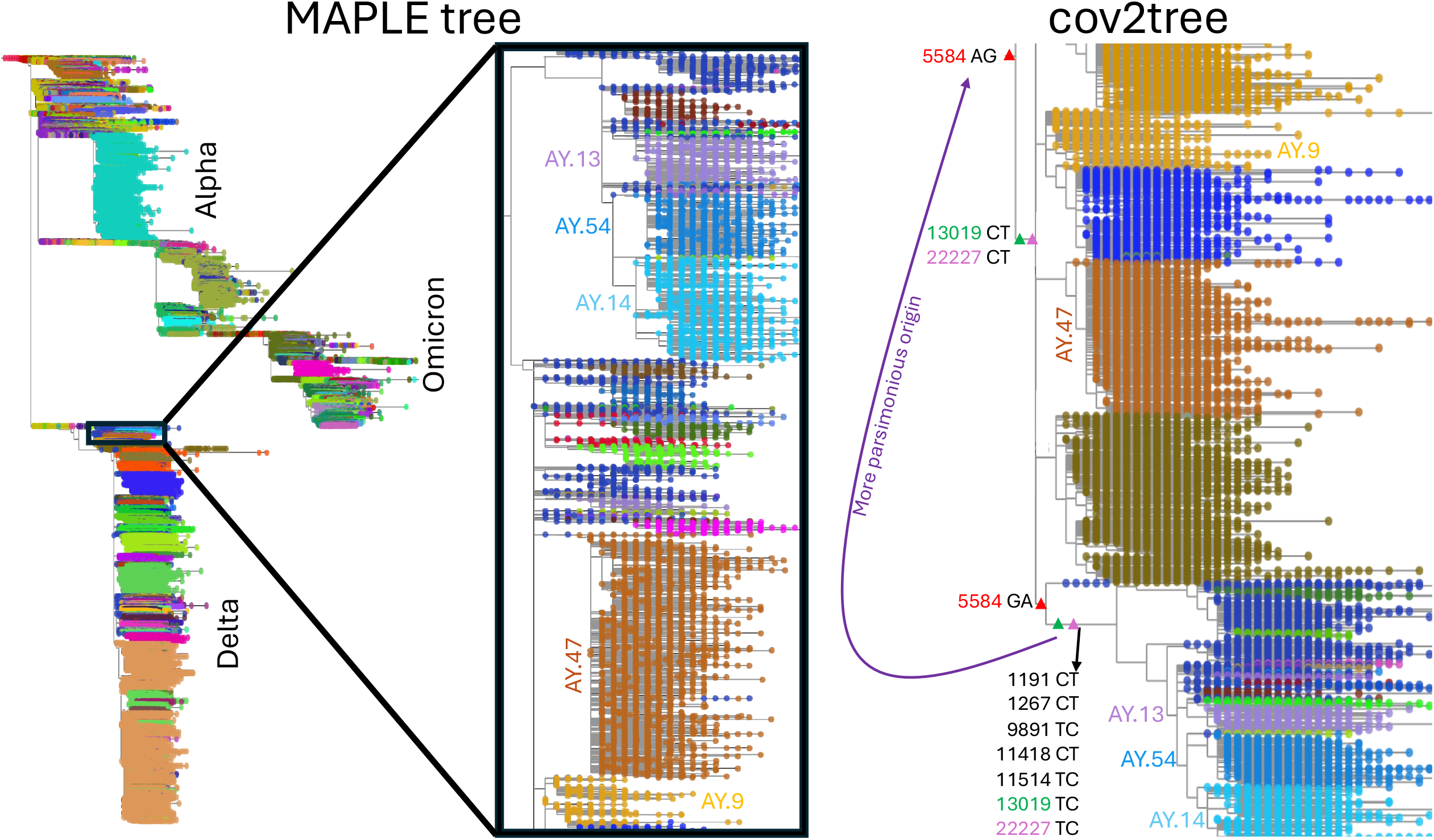
Comparison of the evolutionary histories of the AY SARS-COV-2 lineage inferred by MAPLE and cov2tree. On the left is the global SARS-CoV-2 phylogenetic tree inferred by MAPLE from Viridian genomes. In the center is a zoom-in on part of the tree showing the early evolution of the AY lineage. Phylogenetic tips are colored according to the Pango[32] lineage assigned by Pangolin[33] V4.3 (with pangolin-data V1.21) to the corresponding genomes. We also show the names of some of these lineages. On the right is the corresponding part of the phylogenetic tree for cov2tree[28, 31]. We use colored triangles to annotate estimated mutation events on the branches of the phylogenetic tree, with mutations at the same position shown in the same color. We represent these mutations with their genome position and nucleotides (e.g. ‘5584 AG’ means ‘nucleotide A mutating into G at position 5584’). Three mutations in upstream branches are reverted downstream leading to the clade containing lineages AY.13, AY.14 and AY.54. The purple arrow shows a possible re-grafting of this clade, leading to a more parsimonious evolutionary history avoiding two of these reversions.

It is also useful to investigate the consistency of Pango[32] lineage assignments (made using Pangolin[33]) in our tree: inconsistent assignments, for example showing multiple independent origins of the same lineage, can reveal phylogenetically uncertain lineage histories where our models might help identify more likely scenarios. An example is given in Figure 6, showing that MAPLE infers two independent origins for genomes in the lineage AY.43.3 defined by mutation C to T at position 19955 within lineage AY.43. MAPLE estimates that C19955T occurred two independent times within AY.43, and in total 380 times across the whole tree. Clustering all AY.43.3 genomes in one clade would require replacing one of the occurrences of C19955T with other rarer mutations (Figure 6).

**Figure 6:**
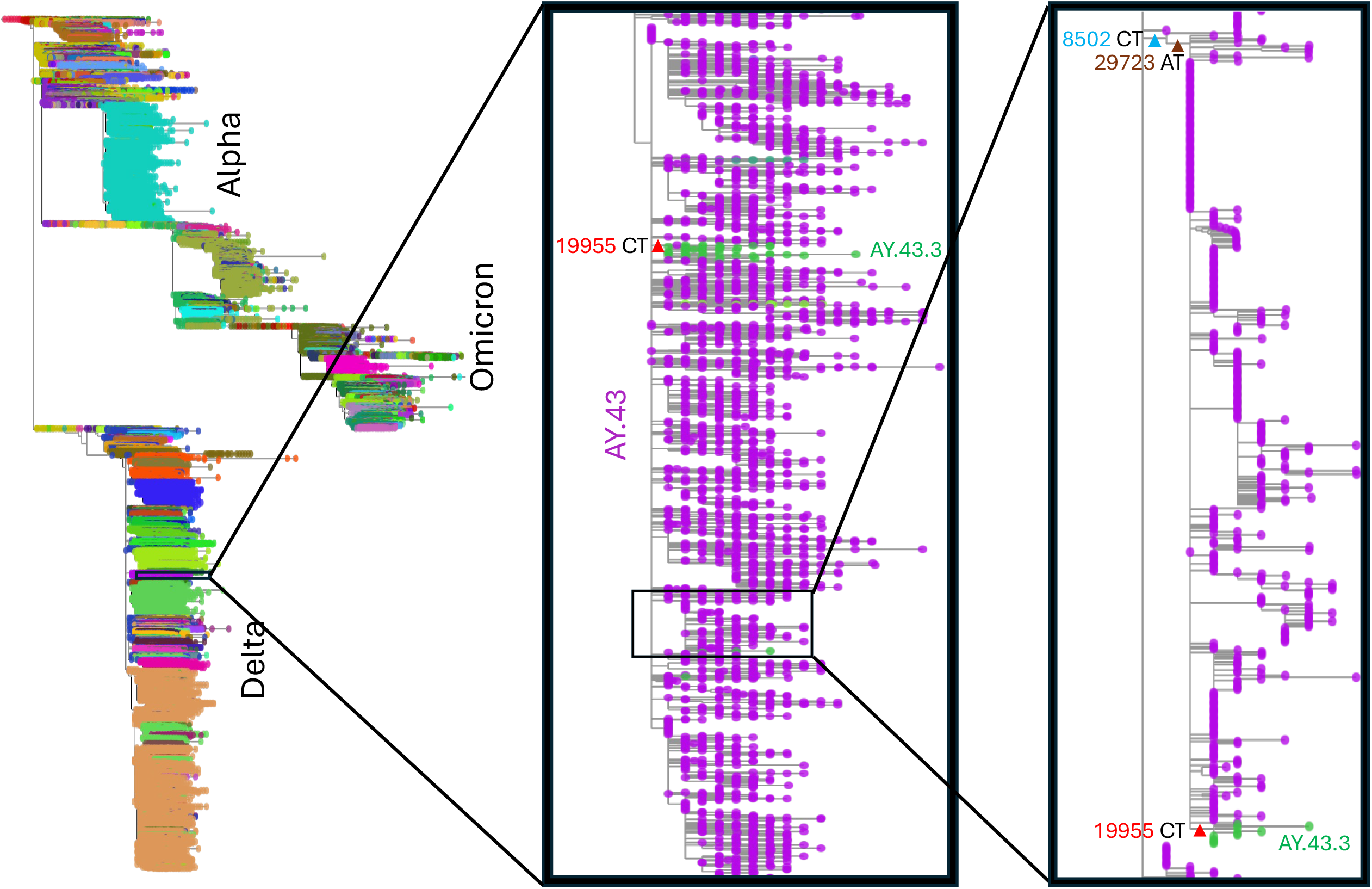
Origin of genomes assigned to lineage AY.43.3. Tree and notation here are as in Figure 5. In the center, we show that the majority of genomes in lineage AY.43.3 are inferred to belong to a clade separated from the rest of the AY.43 genomes by a CT mutation at position 19955. However, in the right panel we show that MAPLE infers a smaller AY.43.3 clade to have originated from a separate 19955 CT mutation. MAPLE infers 19955 CT to be higher frequency mutation (380 separate occurrences in the tree) than the other two mutations (8502 CT, 83 occurrences, and 29723 AT, 33 occurrences) one would need to duplicate to cluster all the AY.43.3 genomes into the same clade.

This approach, however, also highlights possible issues in our tree and alignment. For example, we infer multiple clades of sublineages BA.1.1 and BA.1.15 within BA.1 (Figure 7). This seems to be due to some of these genomes having the consensus sequences of rare recombinants. Investigating read data, we see low coverage and heterozygosity at these putative recombination sites; altogether, these are the hallmarks of consensus sequence artefacts caused by contamination or mixed infections. It seems to us that phylogenetic inference could be improved by better removal of possible contaminated samples. However, with our pre-processing, we already filtered out the majority of available genomes (around 3M out of around 5M), so using more strict filters might be counter-productive. We consider other possible approaches to address these issues in the Discussion section.

**Figure 7:**
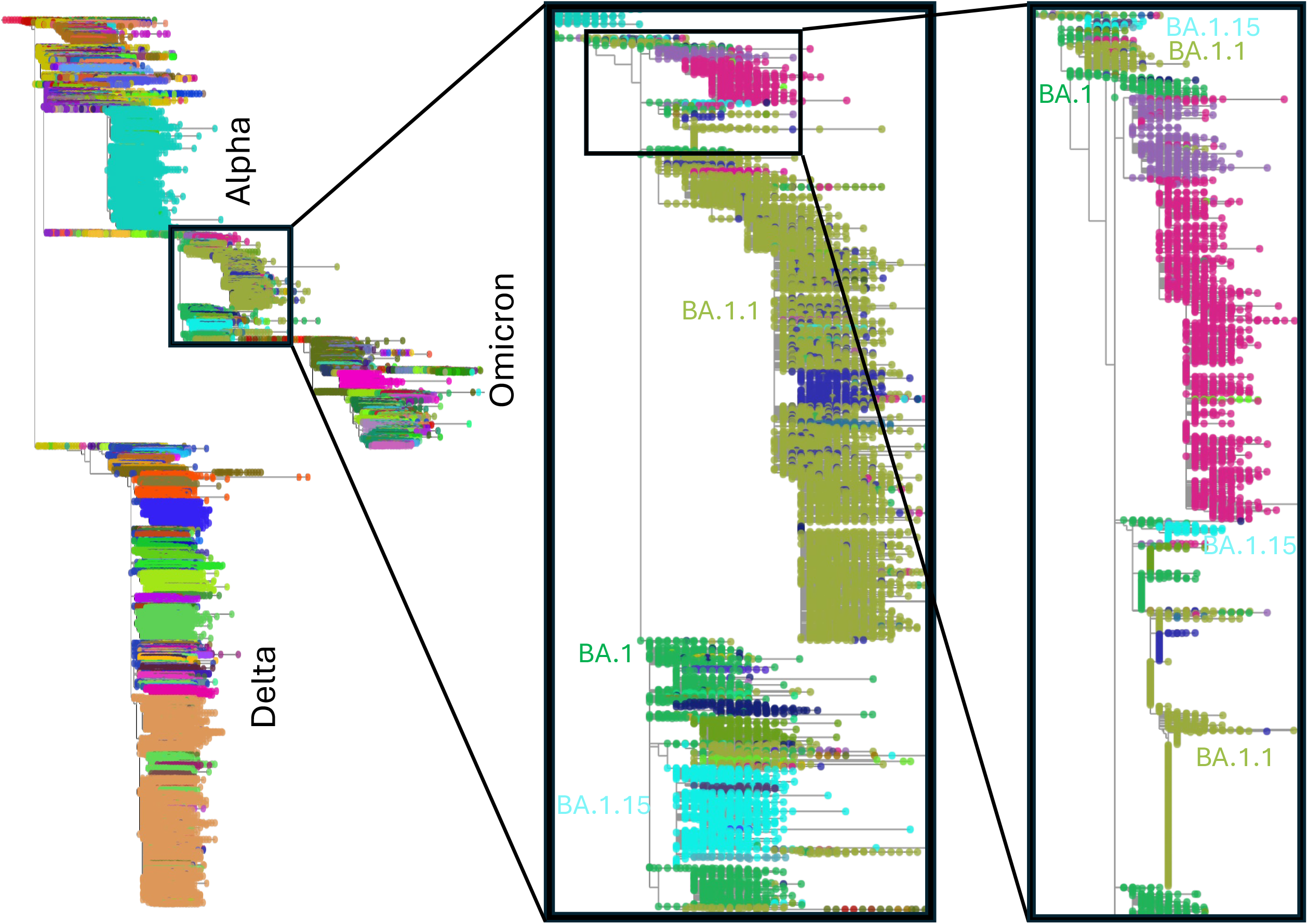
Issues with inferring the history of lineages BA.1.1 and BA.1.15. Tree and notation here are as in Figure 5. As shown in the zoom-ins, lineages BA.1.1 and BA.1.15 appear to have emerged multiple times within parent lineage BA.1, in what we might call ‘duplicate’ clades. Upon investigation, this seems to be caused by possible artefacts in a few consensus sequences, such as SRR18978380 and ERR8227846, due to contamination. Most of the samples in these smaller clades have very low coverage and ambiguous consensus sequence at the gnomic loci of the mutations distinguishing the duplicate clades, meaning that they could equivalently be placed in more than one of these clades.

## 3 Discussion

We introduced a pandemic-scale phylogenetic framework to account for, and estimate, rate variation and recurrent sequence errors. This approach, implemented within the software MAPLE, can lead to improved phylogenetic inference at limited additional computational demand, and allowed us to efficiently investigate recurrent consensus sequence errors. We find that recurrent errors inferred by MAPLE are typically corroborated by sequencing read data. We also implemented algorithmic improvements to MAPLE and parallelized its most computationally demanding tasks. We use these methods to reconstruct a reliable, global, public alignment and phylogenetic tree from *>* 2 million SARS-CoV-2 genomes. These advances are useful not only in studying SARS-CoV-2 evolution and spread, but can be more generally applied within genomic epidemiology, and therefore enhance our preparedness to future pandemics.

There are of course some limitations to our methods. One is the risk of over-parameterization; to prevent this it is important to analyze sufficiently many sequences (we recommend approximately *>* 50, 000 genomes) when considering these models of rate variation and recurrent sequence errors. Furthermore, while MAPLE’s approximations are essential in reducing computational demand and allowing pandemic-scale likelihood-based phylogenetics, they can also lead to biases at higher evolutionary distances [12]. In particular, very high sequence error probability at a given position can lead MAPLE to phylogenetically cluster sequences sharing the same error, interpreting these errors as the result of real mutation events, and therefore underestimating the number of sequence error at the considered position. We observed this phenomenon particularly at positions 8835 and 15521 which have putative consensus sequence errors in hundreds of thousands of genomes. Similar issues can arise when other assumptions of our model are violated, for example if errors at different positions are correlated, or if errors are phylogenetically clustered.

While we showed that these advances can lead to more realistic phylogenetic inference, we also have to consider the extant issues in our alignment and tree. The most prominent of these seems to be due to a small number of genomes whose consensus sequence is seemingly affected by contamination. Only a few of such genomes can noticeably affect the inferred evolutionary history of major clades. While we tried to filter out samples potentially affected by contamination, further improvements with this respect are clearly warranted, and an option might be to consider model-based inference of contamination in read data. Alternatively, stricter consensus calling thresholds could be used, for example more strictly masking regions with relatively low depth or high heterozygosity. However, a probably more appealing avenue could be using a phylogenetic approach, which would not require the use of read data, to identify putative recombinant and contaminated genomes (consensus genomes affected by contamination would not be likely distinguishable from true recombinants with small numbers of descendants), see e.g. [34]. Additionally, including time data into pandemic-scale phylogenetic inference (see e.g. [35]) could improve inference, for example helping us place genomes with substantial sequence uncertainty into more likely regions of the phylogenetic tree.

Implementation of these algorithms and models in C++ is ongoing[36], and will substantially improve their efficiency. We plan to employ OpenMP[37] to implement a shared-memory parallel tree search and significantly reduce memory usage. We will automatically infer the optimal number of cores for parallelizing tree search. The use of multiple local references is also be key to generalize our approach to arbitrary phylogenetic scales, for example switching between parsimony-like approximations and standard phylogenetic likelihood techniques across different branches of the tree, depending on their length.

In conclusion, we presented advances that allow phylogenetic analysis of genomic epidemiological data at unprecedented scale and accuracy, which will be key in tracking transmission and pathogen evolution in a world where genome sequencing plays an ever more relevant role in combating infectious disease.

## 4 Acknowledgements

We thank Russell Corbett-Detig, Yatish Turakhia, Colman O’Cathail, and Chris Bielow for the help in the development of MAPLE and for the useful discussions on these topics. We thank the French Embassy in the UK for funding Z.G. and A.S.

## 5 Methods

### 5.1 Genome lists

We give here a summary of how genome sequences and ancestral likelihoods are stored in MAPLE, which will be relevant to describe many of the method developments presented here. For a full description of the MAPLE approach see [12].

We represent genome sequences and ancestral sequence likelihood in terms of differences with respect to a reference - either local or global. Instead of likelihood vectors, we use genome lists. An entry of a genome list represents the state of a portion of the genome, either a single site, or a collection of contiguous sites. An entry of a genome list is defined as a tuple (*τ, s, l, v*) containing:

- an entry type *τ* ; the allowed types are ‘**R**’, to represent a stretch of contiguous sites identical to the reference; type ‘**N**’ to represent a stretch of contiguous sites containing no descendant sequence information, that is with the four nucleotides having the same partial likelihoods; entries of type **R** and **N** are essential for concisely representing observed and ancestral genomes. Type ‘ **A**’, ‘**C**’, ‘**G**’ and ‘**T**’ to represent a single site where the represented nucleotide is the state of the considered (possibly ancestral) genome with negligeable uncertainty. Type ‘**O**’ is used for a single site where multiple nucleotides have non-negligible partial likelihoods, that is, type **O** entries are used to represent uncertainty at the few genome positions and tree nodes where multiple states are plausible.
- an entry position *s* representing the site(s) of the genome the entry refers to. If the entry represents a stretch of sites, this element is the last one (from 5^*′*^ to 3^*′*^) of these sites. In this case, the first site of the stretch can be inferred by the *s* element of the previous entry in the genome list, since entries in the genome list are sorted according to *s* and since they form a partition of the considered genome.
- the branch length *l* represents the evolutionary distance between the considered node (which stores the genome list) and the location in the tree that the partial likelihoods contained or represented by the genome list entry refer to. For example, an entry of type **R** might represent a stretch of nucleotides observed at the child *c* of the considered node *n*, in case the other child node of *n* has type **N** at these same positions. In this example *l* represents the branch length separating *n* and *c*.
- a vector of relative partial likelihoods *v*, only present in genome list entries of type **O**. The difference between this vector and a standard likelihood vector in phylogenetics is that *v* is normalised (its entries sum up to 1). This is because, while performing tree search, MAPLE only utilizes and stores relative likelihoods, not absolute ones.

### 5.2 Modeling rate variation

Variation in evolutionary rate along the genome can heavily impact phylogenetic inference[16]. As such, models of evolutionary rate variation[18] have been an essential part of phylogenetic analyses in the last decades. Here we first review the most popular phylogenetic models of rate variation (Section 5.2.1) and then present the details of our approach (Section 5.2.2).

#### 5.2.1 Γ random variable model of rate variation

In likelihood-based phylogenetics, variation in evolutionary rates is typically modeled using Gamma-distributed random variable models, where each genome position is allowed to evolve under any of a certain number (typically four) of rates. Under this model, the likelihood is integrated over every possible rate assignment for every site, and a discretized Γ distribution is used to model the distribution of rates across categories[17].

More in detail, for any site *s* of the alignment, for any rate category *c*, and for any node *n* of the tree, one calculates a set of likelihoods 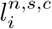, one for each nucleotide *i*; to do this, the substitution rate from nucleotide *i* to nucleotide *j* for category *c* is defined as

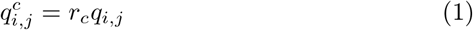

where *r*_*c*_ is the rate of category *c*, and *q*_*i,j*_ is the background substitution rate from *i* to *j*, which is shared between categories. Likelihoods 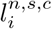 are initialised as usual for each terminal node, site, and category. Then using the Felsenstein pruning algorithm [38] and traversing the child nodes before the parent nodes, one calculates likelihoods at internal nodes, where the likelihoods of class *c*, at each node and site, are calculated using the substitution rates 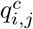 of category *c* and the likelihoods of category *c* from the children nodes. This means that normally the time and memory demands of this approach are *C* times higher than the pruning algorithm without rate variation, with *C* the number of categories considered.

A major problem when trying to use this model with pandemic-scale datasets and algorithms is that, under this model, mutation events in one area of the tree can impact the relative likelihoods of different rate categories in other distant parts of the tree. This is not compatible with the algorithms and data structures in MAPLE, which are based on assumptions of ‘locality’: changes in the tree typically only affect relative likelihoods in a small phylogenetic neighbourhood of the affected nodes. For these reasons, here we used a different model of rate variation, detailed in Section 5.2.2.

#### 5.2.2 Site-specific model of rate variation in MAPLE

Instead of the traditional Γ model of rate variation, we propose a model similar to CAT[19]. In CAT, alignment sites are grouped into a number of classes, and each class of sites has one (and only one) rate assigned to it; this speeds up likelihood calculations since for each site only one set of likelihoods needs to be computed. In CAT, sites are grouped into a small number of classes to reduce the number of rate parameters to avoid overparameterization[19]. In the case of pandemic-scale phylogenetics, since we have millions of observations for each site, we argue that grouping sites into classes is more likely to lead to underparameterization, and we assign to each site *s* its own specific substitution rate *r*_*s*_. Site rates *r*_*s*_ are estimated in MAPLE using Expectation Maximisation (see Section 5.4).

The most important advantage of this site-specific model compared to the Γ one (Section 5.2.1) is that it can be implemented in MAPLE almost without affecting its computational demand. The most important difference is that now, to calculate the likelihood cost of no mutation happening on a stretch of the genome identical to the reference (see [12]), we pre-compute the sums of site-specific substitution rates of the reference nucleotides for prefix segments of the genome.

Another advantage of our approach is that it allows us to estimate biologically informative site-specific rates (see Figure 4). Rates inferred from our 2M genomes Viridian alignment have been uploaded on Zenodo [30].

### 5.3 Sequence error model

Recurrent sequence errors adversely impact phylogenetic inference, especially with low divergence datasets such as collections of SARS-CoV-2 genomes [14, 15]. We previously developed methods to identify and mask recurrent errors [14, 15, 23] to help prevent biases in phylogenetic and other downstream analyses[23]. These approaches have however proved untenable in the long term due to excessive computational demand and the elevated number of genome positions affected by recurrent errors.

Here, we use a different approach. Firstly, we focus on genomes with publicly shared read data, which allows us to consistently run a tailored, state of the art pipeline for calling consensus genomes (Section 5.6.1); this prevents reference-biased calls and therefore reduces the number of recurrent errors in our alignment. The availability of read data also allows us to corroborate putative consensus sequence errors. Secondly, we perform phylogenetic and sequence error inference jointly, using a new recurrent sequence error model within MAPLE, which is detailed in this section. Compared to previous methods, this approach has several advantages:

- It allows the estimation of site-specific error probabilities across the genome and the inference of individual sequence errors.
- It can be paired with a rate variation model (Section 5.2) to disentangle the effects of substitution rate variation and sequence error probability variation.
- Model parameter and sequence error estimation can be performed at the same time as phylogenetic inference.
- It can be used on large datasets of millions of genomes, since it adds limited burden to MAPLE’s computational demand.

While traditional phylogenetic models of sequence errors often assume an homogeneous error probability across the genome[21, 22], we use site-specific error probabilities (see also [39]). Like previous phylogenetic error models, our approach modifies the likelihoods of terminal nodes. For example, without errors, the partial likelihood vector corresponding to an observed nucleotide ‘A’ is defined as (1, 0, 0, 0), where we use the alphabetical order of nucleotides. When accounting for sequence errors with a homogeneous error probability *ϵ*, the same likelihood vector becomes (1 − *ϵ, ϵ/*3, *ϵ/*3, *ϵ/*3)[21, 22]. With site-specific error probabilities, each site *s* of the genome has its own error probability *ϵ*_*s*_, so that, for site *s*, the partial likelihood vector becomes (1 − *ϵ*_*s*_, *ϵ*_*s*_*/*3, *ϵ*_*s*_*/*3, *ϵ*_*s*_*/*3).

Often, modeling site-specific error probabilities with one free parameter per site would result in over-parameterization. However, just like for the substitution rates (Section 5.2.2), the large numbers of sequences we consider compensates for the large number of parameters and allow us to avoid underparameterization, since error probabilities in SARS-CoV-2 are likely highly site-specific[14, 15]. We estimate error probabilities at the same time as rate variation using Expectation Maximisation (Section 5.4).

To efficiently implement the site-specific error model we modify some of the algorithms and data structures in MAPLE. First, we add a flag to each entry of genome lists (which are equivalent to likelihood vectors); a positive flag means that the data/likelihoods represented by the entry refer to a terminal node, and therefore need to be corrected by the probability of a sequence error. Otherwise, a negative flag means that no correction for possible sequence errors is required. This flag does not simply distinguish terminal node entries from internal node entries; this is because the observation of a nucleotide in MAPLE can be carried over from a terminal node likelihood to its parent likelihood, in case the sibling node contains no information at the considered site. In this case, the parent node genome list entry will also have a positive flag. We use flags instead of calculating likelihoods explicitly under the error model because we want to avoid calculation of likelihoods for long stretches of the genome.

When creating new genome lists, or when comparing genome lists to calculate likelihood costs, we need to account for possible sequence errors only when at least one of the two flags of the two genome list entries considered is positive. To do this, we proceed as typical in phylogenetic sequence error models[21, 22], except that, using the assumption of short branches and low error probabilities, we use a first order approximation and ignore terms that are quadratic (or higher order) in either the branch length or the error probability. For example, given a branch length *t*, we ignore terms that are in the order of magnitude of *t*^2^, 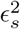, or *t · ϵ*_*s*_. A substantial benefit of this approximation is that we can compute likelihood costs and relative likelihoods for long stretches of the genome in constant time. For example, when two considered genome list entries are of type **R** (they represent a stretch of the genome identical to the reference), then their parent node will also be of type **R** at the same positions, since any alternative would require either 2 mutations, 2 errors, or one error and one mutation at the same position; all these alternative scenarios have negligible probability compared to the history with no mutations and no errors. Conversely, when comparing two genome list entries of type **R** to calculate a likelihood score (for example for calculating the likelihood cost of a placement), we can integrate the cost of no mutations and no errors happening on the considered stretch of the genome in constant time by using pre-computed total sums of error probabilities for genome prefixes, in a very similar way as for accounting for rate variation in MAPLE (Section 5.2.2 and [12]).

### 5.4 Estimation of model parameters with Expectation Maximization

Early versions of MAPLE used a simple, heuristic approach to substitution rate parameter estimation. We now instead use Expectation Maximization to jointly estimate substitution rates, including site-specific ones, and error probabilities, if required. Expectation Maximization parameter optimization is performed after the estimation of the initial tree, and after each round of SPR tree topology improvement. After initial tree estimation, parameter optimization is started from an empirical global substitution rate matrix, uniform site-specific rates, and a uniform site-specific error probability equal to the reverse of the genome length (equivalent to one expected error per genome).

Our Expectation Maximization approach is inspired by Klosterman et al [40] (of which hereby we follow the notation), with the difference that we exploit the shortness of the considered tree branches to reduce computational demand. We first detail our Expectation Maximization approach for the basic scenario that sequence errors and rate variation are not modeled, and then describe the case with sequence errors in Section 5.4.1, and finally we describe the modifications needed to model site-specific rates and error probabilities in Section 5.4.2.

Given a branch of length *T*, and given two nucleotide assignments, *a* for the top of the branch and *b* (with possibly *b* = *a*) for the bottom, we define the substitution probability of such an assignment as *M*_*ab*_(*T*). Assuming *T* ≪ 1 we approximate *M*_*ab*_(*T*) ≈ *Tq*_*ab*_ if *a≠ b* and *M*_*ab*_(*T*) ≈ 1 + *Tq*_*aa*_ otherwise. The expected number of substitutions from nucleotide *i* to *j≠ i* on the considered branch is

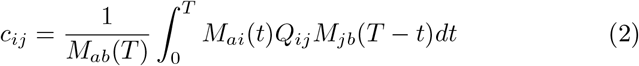

while the expected waiting time in nucleotide *i* on the same branch is

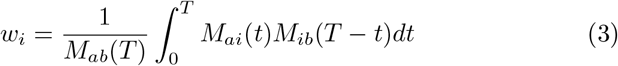

(see [40] for more details). Following again the assumption of short branches, we use first order approximations:

- In case *a* = *b*, we have *w*_*a*_ ≈ *T* and *w*_*i*_ ≈ 0 for *i≠ a*. We also have *c*_*ij*_ ≈ 0 for any *i≠ j*.
- In case *a ≠ b* we have *w*_*i*_ ≈ *T/*2 for *i* = *a* or *i* = *b*, and *w*_*i*_ ≈ 0 otherwise. We also have *c*_*ij*_ ≈ 1 if *i* = *a* and *j* = *b*, and *c*_*ij*_ ≈ 0 otherwise.

*c*_*ij*_ and *w*_*i*_ can be calculated efficiently across genome lists, similarly to how likelihood costs are calculated in MAPLE. Then, the Expectation Maximization estimates of the substitution rates are calculated by traversing the tree and adding up, for each branch encountered, for each genome position, and for each pair of nucleotide assignments *a, b* for the two ends of the considered branch, the value *p*_*ab*_*c*_*ij*_ to a total count *C*_*ij*_ for each nucleotide pair *i, j*, and the value *p*_*ab*_*w*_*i*_ to a total count *W*_*i*_ for each nucleotide *i*, where *p*_*ab*_ is the posterior probability of the assignment *a, b* for the considered branch. These updates are done in constant time for each genome list entry, no matter the number of positions represented by the entry. In case the site-specific model of rate variation (Section 5.2.2) is used, we estimate one rate for each genome position by keeping track of site-specific substitution counts and waiting times. Once we complete the traversal of the tree, the new estimate of the substitution rate *q*_*ij*_ will be *q*_*ij*_ = *C*_*ij*_*/W*_*i*_.

#### 5.4.1 Expectation Maximization with a sequence error model

When we model recurrent sequence errors, as described in Section 5.3, the estimation of model parameters with Expectation Maximization becomes more complex. In addition to branch-specific expectations *c*_*ij*_ and *w*_*i*_, and global counts *C*_*ij*_ and *W*_*i*_, we also track branch-specific expected errors *e*_*ij*_ and global error counts *E*_*ij*_. Even though we do not consider nucleotide-specific error probabilities, we keep the *i, j* subscript for consistency and for possible future extensions. Here we assume, for simplicity, constant substitution rates and error probability along the genome. The needed modifications to model site variation are discussed in Section 5.4.2.

Expectation Maximization counts are calculated as:

- In case *a* = *b* we have *w*_*i*_ ≈ *T* for *i* = *a* and *w*_*i*_ ≈ 0 otherwise. We also have *c*_*ij*_ ≈ 0 and *e*_*ij*_ ≈ 0 for any *i≠ j*.
- In case *a≠ b* and *b* does not derive from a terminal node observation (that is, it could not be the result of a sequence error), we have *w*_*i*_ ≈ *T/*2 for *i* = *a* or *i* = *b*, and *w*_*i*_ ≈ 0 otherwise. We also have *c*_*ij*_ ≈ 1 if *i* = *a* and *j* = *b*, and *c*_*ij*_ ≈ 0 otherwise. *e*_*ij*_ ≈ 0 for any *i≠ j*.
- In case *a≠ b* and *b* could be the result of a sequence error, we have 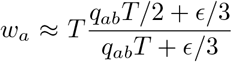 and 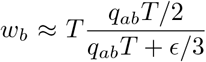, while *w*_*i*_ ≈ 0 if *i≠ a*nd *i≠ b*. We also have 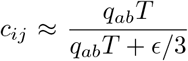 if *i* = *a* and *j* = *b*, and *c*_*ij*_ ≈ 0 otherwise. Finally, 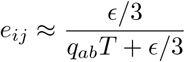 for *i* = *a* and *j* = *b*, and *e*_*ij*_ ≈ 0 otherwise.

We traverse the tree and add up, for each branch encountered and each pair of nucleotide assignments *a, b* for the two ends of the considered branch, the value *p*_*ab*_*c*_*ij*_ to the total count *C*_*ij*_ for each *i, j*, the value *p*_*ab*_*w*_*i*_ to the total count *W*_*i*_ for each *i*, and the value *p*_*ab*_*e*_*ij*_ to the total count *E*_*ij*_ for each *i, j*. Again, these updates are done in constant time for each genome list entry (possibly covering thousands of genome positions) considered. Once we complete the traversal of the tree, the new estimate of the substitution rate *q*_*ij*_ is *q*_*ij*_ = *C*_*ij*_*/w*_*i*_. The new estimate of the error probability is *ϵ* = Σ_*i,j*_ *E*_*ij*_*/K*, where *K* is the number of times informative characters (non-’N’ characters) are observed at terminal nodes (we do not count *w*_*i*_ on branches/positions above type ‘N’ characters).

#### 5.4.2 Expectation Maximization with site variation

When modeling site-specific rates and error probabilities, we need to keep separate counts for each genome positions, that is, instead of *W*_*i*_ we define total waiting time for character *i* and site *s* as 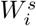; we add to this count only the waiting times *w*_*i*_ calculated at position *s* of the alignment. We similarly define and update position-specific counts 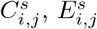, and *K*^*s*^.

### 5.5 Other new features

#### 5.5.1 Online inference: partial initial tree and sample placement

We have implemented in MAPLE the option of using an initial tree, either complete (relating all the sequences in the input alignment) or incomplete. If the initial tree is incomplete, MAPLE will add the missing samples to the tree using maximum likelihood stepwise addition, and then perform SPR searches to improve the tree topology. We have also implemented the option to perform a faster, partial SPR search targeting only nodes whose genome lists have been affected during stepwise addition. If only a very few samples have been added to the initial tree, this partial SPR search can be very fast.

These features not only allow users to specify custom initial trees, but they also allow online phylogenetic inference[41] (where new samples are added to a tree as they become available, avoiding costly repeated reestimation of a large phylogeny from scratch) and placement of individual samples onto a target tree[10].

#### 5.5.2 Improvements to the placement and SPR search algorithms

In earlier versions of MAPLE, placement and SPR searches were divided in two stages. In the first stage we traversed the tree to find a preliminary optimal placement. Then, in the second stage, we optimised the exact location of the placement and the new branch lengths near the initial optimal placement[12].

We have now implemented more thorough searches, similarly divided in two stages, but in which the first stage returns a set of preliminary near-optimal placements (instead of just one). The second stage then optimises branch lengths for all these candidates, and selects the best placement after this optimisation. This improves tree inference accuracy at very limited additional computational demand.

#### 5.5.3 Tree rooting

We implemented an efficient dynamic programming search of the maximum likelihood tree root location. This algorithm traverses the tree from parents (starting from the current tree root) to children nodes. As we traverse nodes, we keep track of how phylogenetic likelihoods are affected by moving the tree root from a parent node to its child, with the consequent inversion of mutation events on the branch separating the two.

#### 5.5.4 Branch length optimization

We have implemented in MAPLE a more efficient and accurate procedure for branch length optimization, based on approximating the derivative of the phylogenetic relative log-likelihood as a function of branch length.

Given a branch of the phylogenetic tree, assume that *G*_2_ is the genome list of the bottom node of the branch (representing the relative likelihoods of the subtree below the considered branch), and *G*_1_ is the genome list for the top node of the branch (representing the relative likelihoods of the subtree complementary to the subtree below the considered branch). Our aim is to efficiently calculate the branch length value *t* that minimizes the likelihood cost of merging *G*_1_ and *G*_2_. This likelihood cost is the product of the likelihood cost of each individual intersection entry of *G*_1_ and *G*_2_ (see [12]), so this is the same as minimizing the sum of the log-likelihoods of each intersection entry. To do this, we look for the values of *t* for which the derivative of the likelihood cost is 0. Assuming that *E*_1_ is the considered entry of *G*_1_ and *E*_2_ is the considered entry of *G*_2_, we can approximate the derivative with respect to *t* of the log-likelihood costs as:

- If *E*_1_ and *E*_2_ represent the same nucleotide, or the same stretch of nucleotides, the log-likelihood cost is ≈ Σ_*i*_ −*r*_*i*_(*t* + *t*_0_) where *r*_*i*_ is the substitution rate of the reference nucleotide at position *i*, and *t*_0_ is the sum of the branch length elements of *E*_1_ and *E*_2_ (these are used in the presence of ‘N’ characters in the alignment). The derivative is then − Σ_*i*_ *r*_*i*_.
- If *E*_1_ and *E*_2_ represent different nucleotides *i* and *j*, the log-likelihood cost is ≈ log(*q*_*ij*_) + log(*t* + *t*_0_), where *q*_*ij*_ is the substitution rate from *i* to *j*. Its derivative is 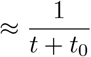.
- If any of *E*_1_ or *E*_2_ is of type N (that is, it provides no sequence information), the log-likelihood cost is null, and so is its derivative.

If any of *E*_1_ or *E*_2_ is of type O (meaning that multiple nucleotides are possible at this site and node), then we have to consider the different relative likelihoods of different nucleotides. Here we consider the most complex case of both *E*_1_ and *E*_2_ being of type O, the other scenarios being simplifications of the following. We call *p*_1_ the relative likelihood vector of *E*_1_ and *p*_2_ the relative likelihood vector of *E*_2_. The log-likelihood cost is ≈ log (Σ_*i*_ *p*_1_(*i*)*p*_2_(*i*) + (*t* + *t*_0_) _*i,j*_ *q*_*ij*_*p*_1_(*i*)*p*_2_(*j*)). We don’t know a priori if *p*_*i*_(*i*) and *p*_2_(*j*) are large or small compared to and. However, if the coefficient of the *t*, Σ_*i,j*_ *q*_*ij*_*p*_1_(*i*)*p*_2_(*j*),, is negative, then we know that Σ_*i*_ *p*_1_(*i*L)*p*_2_(*i*) is the dominant term inside the logarithm. In fact, (*t* + *t*_0_) Σ_*i,j*_ *q*_*ij*_*p*_1_(*i*)*p*_2_(*j*) is equal to (*t* + *t*_0_) Σ_*i*≠*j*_ *q*_*ij*_*p*_1_(*i*)*p*_2_(*j*) + (*t* + *t*_0_) Σ_*i*_ *q*_*ii*_*p*_1_(*i*)*p*_2_(*i*) of which the left side is positive and the right side is negative (and therefore larger in absolute values since we assumed that the sum of the two terms is negative); since we assume that none of the substitution rates *q*_*ii*_ is very large (that is, it’s not in the order of magnitude of the reverse of a typical branch length, which would break the assumptions of MAPLE), then Σ_*i*_ *q*_*ii*_*p*_1_(*i*)*p*_2_(*i*) and therefore Σ_*i,j*_ *q*_*ij*_*p*_1_(*i*)*p*_2_(*j*) is in the same or smaller order of magnitude than Σ_*i*_ *p*_1_(*i*)*p*_2_(*i*), so we obtain |(*t* + *t*_0_) Σ_*i,j*_ *q*_*ij*_*p*_1_(*i*)*p*_2_(*j*)| ≪ *Σ*_*i*_ *p*_1_(*i*)*p*_2_(*i*). By expressing the log-likelihood cost as 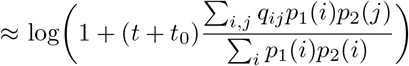 we can then approximate it as 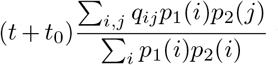 which has derivative with respect to *t* of 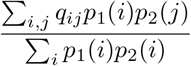. We cannot necessarily use this approximation when Σ_*i,j*_ *q*_*ij*_*p*_1_(*i*)*p*_2_(*j*) *>* 0; instead, in this case we can express the log-likelihood cost as *c* + log(*t* + *a*) for some positive values *a* and *c*, whose derivative is therefore 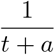.

In all cases, the derivative of the approximate log-likelihood cost of an entry is in one of the two following forms:

- −*r* for some positive cumulative substitution rate *r*.
- 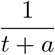 for some positive value *a*.

Summing across all entry intersections, the total approximate log-likelihood derivative becomes 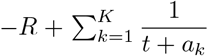, for some positive *R, K*, and *a* values. This expression is well defined for *t >* 0 and is strictly decreasing as *t* increases, with limit −*R* at *t* → +∞ and limit +∞ at either *t* = 0 or some value below 0. This means that either there is only one *t >* 0 for which this derivative is 0, and therefore 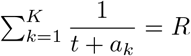, which then corresponds to the optimal branch length, or the optimal branch length is *t* = 0. To find in practice the value of *t* for which 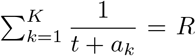 we use a bisection algorithm. Initial upper and lower bounds for the bisection algorithm are set to *t* = *K/R* − max_*k*_(*a*_*k*_) and *t* = *K/R* − min_*k*_(*a*_*k*_), with exceptions in case any of these values is negative.

#### 5.5.5 Robinson-Foulds distances

We have implemented an efficient Robinson-Foulds distance[42] calculation in MAPLE for large phylogenetic trees. We use the Day 1985 [43] linear time and memory algorithm. Note that other approaches might be marginally more efficient (see e.g. fastRF [44] and Hush-RF [45]), but our implementation is particularly useful when comparing trees with short divergence. This is because we distinguish between branches with positive length and branches with 0 length. We consider a branch of length 0 as non-informative, that is, the clade defined by the branch is not considered part of the tree topology. Our implementation allows the specification of a minimum branch length threshold under which branches are considered absent from the tree (effectively of length 0); this helps preventing biases in tree distance calculations when considering trees that are effectively multifurcating, even though they might be represented as binary, as usually is the case for the output trees of popular phylogenetic methods[46–48].

### 5.6 SARS-CoV-2 genome datasets

#### 5.6.1 Viridian and GenBank SARS-CoV-2 genome dataset

To investigate possible recurrent errors in different consensus sequence calling pipelines, we created a dataset of genomes for which we could find read data and consensus sequences both from Viridian and GenBank. The approach to data collection is described in[27]. We separated the Viridian and GenBenak consensus sequences to create two separate alignments. The genomes were aligned to the reference MN908947.3 using MAFFT v7.505 with options ‘–auto –keeplength –addfragment’. We then filtered out genomes with either:

1. 3000 or more ‘N’ or ‘-’ characters in either consensus sequence,
2. 50 or more blocks of consecutive ‘N’ or ‘-’ characters in either consensus sequence,
3. 5 or more IUPAC ambiguity characters in either consensus sequence, since these genomes might have been affected by low coverage or contamination. This resulted in 2 alignments each with 2,993,121 consensus genomes one containing Viridian consensus sequences, and the other GenBank consensus sequences for the same genomes. We then ran multiple times MAPLE 0.6.7 with error model on each alignment, iteratively masking positions that appeared most strongly affected by recurrent sequence errors (see Section 2.4).

After this, we created a second alignment containing all the high quality Viridian genomes available to us, and masking initial (1-78) and final (29769-29903) positions, and alignment columns affected by putative recurrent errors (Table 2.4.1). Like before, we removed genomes that might have been affected by contamination, mixed infections and low coverage, this time with more stringent thresholds:

1. 2500 or more ‘N’ or ‘-’ characters,
2. 30 or more blocks of ‘N’ or ‘-’ characters,
3. 4 or more IUPAC ambiguity characters.

We also filtered out genomes based on coverage and heterozygosity patterns in the read data, that is, we removed genomes with either:

1. *>* 2500 positions with coverage ≤ 100,
2. *>* 1500 positions with coverage ≤ 20,
3. *>* 30 positions with *>* 5% minor allele frequency,
4. *>* 7 positions with *>* 10% minor allele frequency,
5. *>* 2 positions with *>* 20% minor allele frequency.

From the resulting alignment, we masked deletions of length ≤ 30 bp, that is, we converted them to the reference sequence. While deletions do not usually cause problems to phylogenetic inference, and are instead usually interpreted as missing data, we observed that, in this case, errors (either in sequence or alignment) in a few genomes at position of common deletions caused wrong substitutions to be inferred on ancestral branches, in addition to causing substantial phylogenetic uncertainty and instability (with consequently longer runtime for phylogenetic inference). We did not mask longer deletions in the consensus genomes as these often represent parts of the genomes with low coverage, and as such do not cluster together phylogenetically and so do not seem to cause these problems.

The resulting global Viridian alignment contains 2,072,111 genomes. From this alignment, we estimated a phylogenetic tree in MAPLE V0.6.8 under an UNREST model with rate variation in three steps:

- First we estimated an initial tree with 1 core, with options ‘—noFastTopologyInitialSearch --numTopologyImprovements 0’.
- Then, starting from the tree inferred in the first step (‘--inputTree’ option), we used 14 cores to improve the tree topology with option ‘--largeUpdate’.
- Finally, starting from the tree inferred in the second step, we ran a deeper run of topology improvement with 14 cores with options ‘--largeUpdate --noFastTopologyInitialSearch –thresholdLogLKtopology 28.0 --allowedFailsTopology 8’.

The alignment, metadata, inferred tree, and inferred substitution rates have been uploaded on Zenodo [30].

#### 5.6.2 Simulated SARS-CoV-2 sequence data

We simulated SARS-CoV-2 genomes evolved according to a known (‘true’) background phylogeny and substitution model. We used as background tree the publicly available 26 October 2021 global SARS-CoV-2 phylogenetic tree from http://hgdownload.soe.ucsc.edu/goldenPath/wuhCor1/UShER_SARS-CoV-2/^31^ representing the evolutionary relationship of 2,250,054 SARS-CoV-2 genomes as obtained using UShER^10^.

We used phastSim v0.0.3^49^ to simulate sequence evolution along this tree according to SARS-CoV-2 non-stationary neutral mutation rates^13^ and using the SARS-CoV-2 Wuhan-Hu-1 genome^50^ as root sequence. We also simulated variation in substitution rates, using in phastSim a gamma distribution of rates with *α* = 0.2, leading to approximately thesave substitution rate variance as in the substitution rates estimated with MAPLE from the GISAID real SARS-CoV-2 genome dataset considered in [12].

Ambiguity characters were introduced in the alignment to mimic sequence incompleteness observed in real SARS-CoV-2 genome data. For each simulated sequence we sampled one random sequence from the GISAID real SARS-CoV-2 genome dataset considered in [12] and copy-pasted from it the stretches of ‘N’ and gap ‘-’ characters into the simulated sequence. Additionally, we counted the number of isolated ambiguous characters in the real sequence, and we masked an equal number of randomly selected SNPs (differences with respect to the reference genome) in the simulated sequence. See [12] for more detail.

Finally, we added sequence errors to the simulated alignment with a custom script; we used the same site-specific sequence error probabilities estimated from the GISAID SARS-CoV-2 genome dataset in [12] using MAPLE with rate variation and sequence error model.

